# Autoantibodies against Progranulin and IL-1 receptor antagonist due to immunogenic posttranslational isoforms contribute to hyperinflammation in critically ill COVID-19

**DOI:** 10.1101/2021.04.23.441188

**Authors:** Lorenz Thurner, Natalie Fadle, Moritz Bewarder, Igor Kos, Evi Regitz, Bernhard Thurner, Yvan Fischer, Onur Cetin, Torben Rixecker, Marie-Christin Hoffmann, Klaus-Dieter Preuss, Claudia Schormann, Frank Neumann, Sylvia Hartmann, Theresa Bock, Dominic Kaddu-Mulindwa, Birgit Bette, Klaus Roemer, Joerg Thomas Bittenbring, Konstantinos Christofyllakis, Angelika Bick, Vadim Lesan, Zanir Abdi, Sebastian Mang, André Becker, Carlos Metz, Frederik Seiler, Johannes Lehmann, Philipp Agne, Thomas Adams, Andreas Link, Christian Werner, Angela Thiel-Bodenstaff, Matthias Reichert, Guy Danziger, Sophie Roth, Cihan Papan, Jan Pilch, Thorsten Pfuhl, Patrick Wuchter, Christian Herr, Stefan Lohse, Hubert Schrezenmeier, Michael Boehm, Frank Langer, Gereon Gäbelein, Bettina Friesenhahn-Ochs, Christoph Kessel, Dirk Foell, Robert Bals, Frank Lammert, Sixten Körper, Jürgen Rissland, Christian Lensch, Stephan Stilgenbauer, Sören L. Becker, Sigrun Smola, Marcin Krawczyk, Philipp M. Lepper

**Author notes:** shared last authorship (Marcin Krawczyk, Philipp M. Lepper). Corresponding author: Lorenz Thurner, MD, Dept. of Internal Medicine I and José-Carreras-Center for Immuno- and Gene Therapy Saarland University Medical School, D-66421 Homburg/Saar, Germany.

## Abstract

Hyperinflammation is frequently observed in patients with severe COVID-19. Inadequate and defective IFN type I responses against SARS-CoV-2, associated with autoantibodies in a proportion of patients, lead to severe courses of disease. In addition, hyperactive responses of the humoral immune system have been described.

In the current study we investigated a possible role of neutralizing autoantibodies against antiinflammatory mediators. Plasma from adult patients with severe and critical COVID-19 was screened by ELISA for antibodies against PGRN, IL-1-Ra, IL-10, IL-18BP, IL-22BP, IL-36-Ra, CD40, IFN-α2, IFN-γ, IFN-ω and serpinB1. Autoantibodies were characterized and the antigens were analyzed for immunogenic alterations.

In a discovery cohort with severe to critical COVID-19 high titers of PGRN-autoantibodies were detected in 11 of 30 (36.7%), and of IL-1-Ra-autoantibodies in 14 of 30 (46.7%) patients. In a validation cohort of 64 patients with critical COVID-19 high-titer PGRN-Abs were detected in 25 (39%) and IL-1-Ra-Abs in 32 of 64 patients (50%). PGRN-Abs and IL-1-Ra-Abs belonged to IgM and several IgG subclasses. In separate cohorts with non-critical COVID-19, PGRN-Abs and IL-1-Ra-Abs were detected in low frequency (i.e. in < 5% of patients) and at low titers. Neither PGRN-nor IL-1-Ra-Abs were found in 40 healthy controls vaccinated against SARS-CoV-2 or 188 unvaccinated healthy controls. PGRN-Abs were not cross-reactive against SARS-CoV-2 structural proteins nor against IL-1-Ra. Plasma levels of both free PGRN and free IL-1-Ra were significantly decreased in autoantibody-positive patients compared to Ab-negative and non-COVID-19 controls. In vitro PGRN-Abs from patients functionally reduced PGRN-dependent inhibition of TNF-α signaling, and IL-1-Ra-Abs from patients reduced IL-1-Ra- or anakinra-dependent inhibition of IL-1ß signaling. The pSer81 hyperphosphorylated PGRN isoform was exclusively detected in patients with high-titer PGRN-Abs; likewise, a hyperphosphorylated IL-1-Ra isoform was only found in patients with high-titer IL-1-Ra-Abs. Thr111 was identified as the hyperphophorylated amino acid of IL-1-Ra. In longitudinally collected samples hyperphosphorylated isoforms of both PGRN and IL-1-Ra emerged transiently, and preceded the appearance of autoantibodies. In hospitalized patients, the presence of IL-1-Ra-Abs or IL-1-Ra-Abs in combination with PGRN-Abs was associated with a higher morbidity and mortality.

To conclude, neutralizing autoantibodies to IL-1-Ra and PGRN occur in a significant portion of patients with critical COVID-19, with a concomitant decrease in circulating free PGRN and IL-1-Ra, indicative of a misdirected, proinflammatory autoimmune response. The break of self-tolerance is likely caused by atypical hyperphosphorylated isoforms of both antigens, whose appearances precede autoantibody induction. Our data suggest that these immunogenic secondary modifications are induced by the SARS-CoV-2-infection itself or the inflammatory environment evoked by the infection and predispose for a critical course of COVID-19.

## INTRODUCTION

COVID-19 caused by SARS-CoV-2 shows a very wide spectrum of manifestations and severity ranging from completely asymptomatic infection, to mild cold symptoms or attenuation of the sense of taste and smell, to acute respiratory distress syndrome (ARDS), the latter often associated with thromboembolic complications (*1*),(*2*),(*3*),(*4*). Patients requiring intensive care treatment often present with a hyperinflammatory state, which has been compared with hemophagocytic lymphohistiocytosis (HLH)(*5*). Accordingly, antiinflammatory or immunosuppressive drugs like dexamethasone or inhibitors of the IL-1ß-, IL-6-, JAK/STAT-, or the BCR-pathways have been studied in clinical trials (*6*),(*7*),(*8*),(*9*),(*10*),(*11*),(*12*). Among these drugs, though, only dexamethasone and IL-6 pathway blockade have so far been identified to provide a significant clinical benefit, in terms of a reduced 28-day mortality in patients with severe COVID-19 (*6*).

COVID-19 has been suggested to resemble hemophagocytic lymphohistiocytosis (HLH) (*5*), a condition associated with a hyperinflammatory dysregulation of the immune system. Misdirected, or defective IFN I responses were reported as a key factor for inefficient immune responses in patients with COVID-19 (*13*),(*14*),(*15*). In this context autoantibodies against IFN I were detected specifically in patients with severe courses of COVID-19 (16). Moreover, exuberant B-cell responses were regularly found in COVID-19 (*17*),(*18*), along with a high prevalence of antiphospholipid- (*19*), anti-Ro/La-(*20*) or anti-annexin-V-antibodies (*21*). A role of proinflammatory antibodies has so far not yet been clearly established (*22*).

Previously, we had identified neutralizing autoantibodies against progranulin (PGRN) in sera from patients with primary small vessel vasculitis (*23*). Subsequently we found progranulin-Abs in various rheumatic and other autoimmune diseases, but only very rarely in healthy controls, elderly or obese subjects, patients with sepsis and patients with melanoma (*24*),(*25*),(*26*).

PGRN, also called proepithelin, is a secreted precursor protein. Beside several other biological functions (*27*), a major property of PGRN is its anti-inflammatory activity (*28*),(*29*), which is mediated by direct binding to TNFR1, TNFR2 and DR3 and thus antagonization of responses to TNF-α and TL1-a (*30*),(*31*). This has been demonstrated in vivo in several mouse models including collagen and collagen-antibody induced arthritis (*30*), OXA induced dermatitis (*32*), and more relevant for COVID-19, also in LPS-induced lung injury/ARDS mouse models (*33*),(*34*),(*35*). Both PGRN and TNF-α bind to cysteine-rich domain 2 and 3 of TNFR (*36*). The proinflammatory effect of neutralizing PGRN-Abs was characterized in vitro by the analysis of TNF-α-induced cytotoxic effects by MTT-assays and by downmodulation of FoxP3 in CD4^+^CD25^hi^ Tregs in inflammatory bowel diseases and rheumatic diseases (*25*),(*37*). With hyperinflammatory states often observed in patients with severe COVID-19, and in light of similarities between this viral condition, vasculitis and autoimmune diseases (*4*), the aim of the current study was to investigate the possible occurrence of antibodies directed against previously described anti-inflammatory antigens against as progranulin (*23*) or IL-10 (*38*), but also against other hypothesis-driven selected secreted anti-inflammatory mediators like interleukin-1 receptor antagonist (IL-1-Ra).

## RESULTS

### Presence, titers and IgG subclasses of anti-progranulin and anti-IL-1-Ra antibodies

The discovery cohort consisted of 30 patients with COVID-19 (25 male, 5 female, median age 60 years, 21 with critical disease requiring treatment in ICU and 9 with moderate to severe disease not in ICU, median plasma ferritin: 1547 ng/mL, median CRP 91.3 mg/L). The anti-PGRN ELISA demonstrated the presence of PGRN-autoantibodies in plasma of 11/30 patients (36.7%), of which 9 were treated in ICU and 10 were males (median ferritin: 2159.5 ng/ml, median CRP: 160.1 mg/l) (Fig. 1 A). In a control-cohort of non-COVID-19 ICU patients, only 1 of 28 patients (3.6%) had weakly detectable PGRN-Abs (Fig. 1 B). In autoantibody-positive patients with severe or critical COVID-19, the titers ranged from 1:1600 up to 1:3200 (Fig. 1 C). PGRN-Abs belonging to the IgM class were found in 10 of 11 patients with COVID-19, and in all of the 11 PGRN-Ab-positive patients IgG class autoantibodies were found, with IgG1 detectable in 8, IgG2 in 9, IgG3 in 6 and IgG4 in 8 patients (Fig. 1 D).

**Figure 1:**
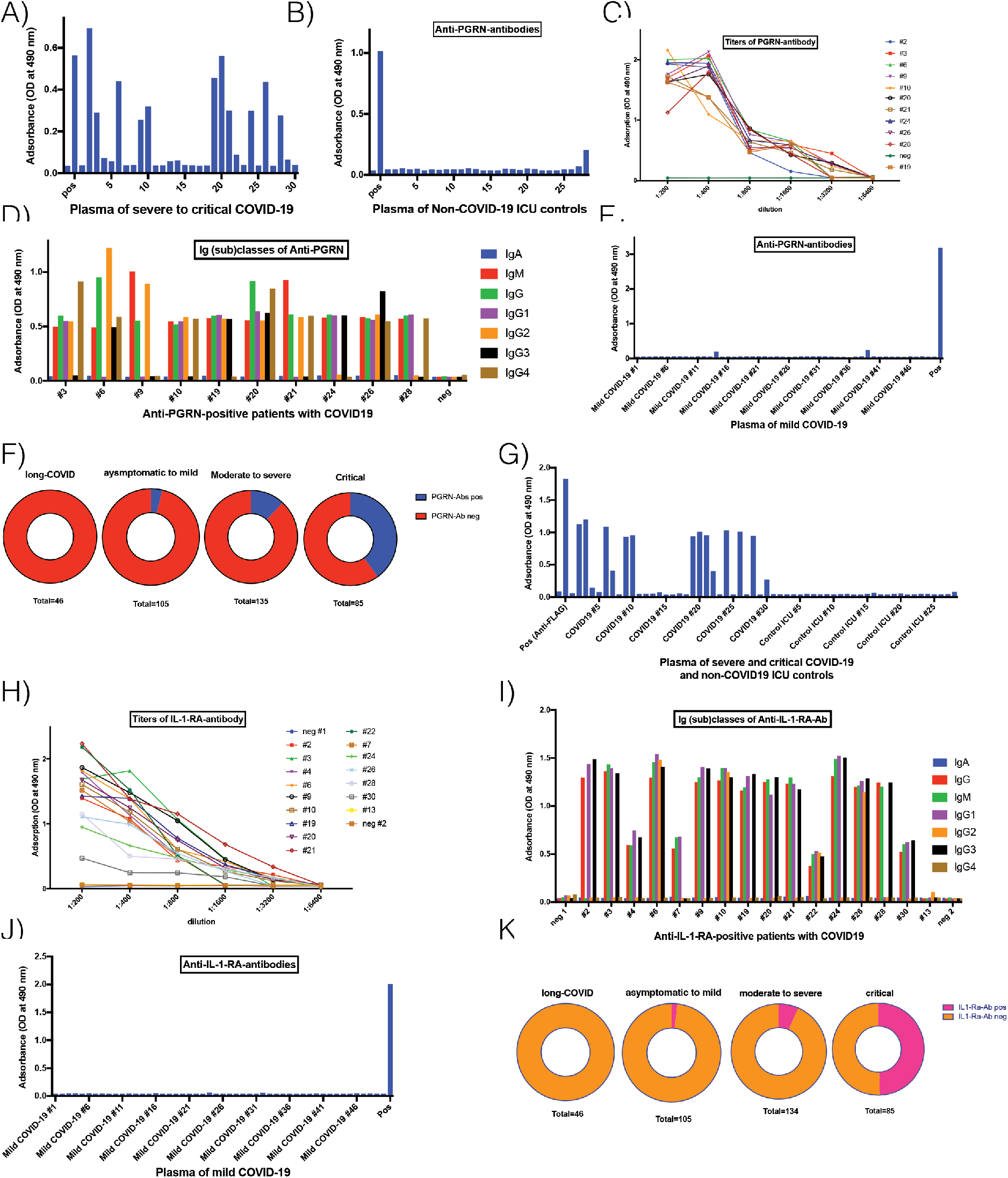
A) Occurrence of PGRN-antibodies in plasma from patients with moderate to severe COVID-19 of the discovery cohort and B) in control patients from intensive care unit without COVID-19. C) Titers of PGRN-Abs from patients with COVID-19 of the discovery cohort. D) Ig classes and IgG subclasses of autoantibodies. E) PGRN-Abs were detected at low titers in two patients with asymptomatic or mild COVID-19. F) Collectively, PGRN-Abs with relevant titers were detected in patients with severe COVID-19 of the discovery and validation cohorts with severe COVID-19 34 of 85 (40%)22 of 65 patients (33.8%), compared to an occurrence of 16 of 135 (11.9%) in moderate COVID-19 and 4 of 105 (3.8%) in mild or asymptomatic COVID-19. G) Occurrence of IL-1RA-Abs in patients with moderate to severe COVID-19 of the discovery cohort and in controls from patients from intensive care unit without COVID-19. H) Titers of IL-1-Ra-Abs from patients with COVID-19 and I) Ig classes and IgG subclasses. J) Only weak, but no relevant IL-1-Ra-Abs were detected in patients with asymptomatic or mild COVID-19. K) Collectively, IL-1-Ra-Abs with relevant titers were detected in 42 of 85 (49,4%) patients with critical severe COVID-19 of the discovery and validation cohorts with severe COVID-19 in 30 of 65 patients (46.2%), compared to an occurrence of 4 9 of 134 (7.6%) hospitalized patients with moderate to severe COVID-19, compared to 2 of 105 (1.9%) in 205 patients (2%) of patients with mild or asymptomatic, mild or moderate COVID-19.

In a validation cohort of 64 patients (median age: 61 years, 53 male, 11 female) with critical COVID-19 requiring mechanical ventilation, 25 of 64 (39%) patients had PGRN-Abs (Supplementary Fig. 1 A). The median age of PGRN-Ab-positive patients was 60 years. Very similar to the discovery cohort, the PGRN-Abs often belonged to both the IgM class and different IgG subclasses and had similar titers (Supplement Fig. 1 B and 1C). In a cohort of 126 hospitalized patients with predominantly moderate COVID-19 (median age: 65.5 years, 58% male, 42% female), PGRN-Abs were only detected in 16 (12.7%) (Supplementary Fig. 1 E and F). Alltogether, 4 of 105 (3.8%) patients with predominantly mild courses of COVID-19 had PGRN-Abs at low titers: PGRN-Abs were present in 2 of 56 (3.5%) convalescent plasma donors from the CAPSID-trial with an asymptomatic to mild course of COVID-19 (supplementary Figure 1 I) and in 2 of 49 patients (4.1%) with mild COVID-19 from the Institute of Virology, Saarland Medical Center (Fig. 1 E).

Grouping the above results according to disease severity: If all critically ill patients with COVID-19 with ICU treatment of the discovery and validation cohort are considered together, 34 of 85 patients (40%) had PGRN-Abs mostly at high titers, compared to 16 of 135 (11.9%) hospitalized patients with moderate to severe COVID-19 (Fisher exact test: *p* < 0.0001) with PGRN-Abs at predominantly low titers, and to 4 of 105 (3.8%) with asymptomatic or mild COVID-19 (Fisher exact test: *p* < 0.0001) at low titers (Figure 1 F). PGRN-Abs were not detected in serum or plasma of 40 healthcare-workers after 2 vaccinations against SARS-CoV2 (Supplementary Fig. 1 I), and PGRN-Abs were also not detected in 75 patients with long-COVID.

Next, we analyzed autoantibodies against further anti-inflammatory mediators. IL-1-Ra-antibodies were detected in 14 of 30 patients of the discovery cohort (46,7%, median age: 60.5 years, 11 male, 3 female, 10 treated on ICU, 4 treated on normal ward, median ferritin: 1961 ng/ml, median CRP: 121.7 mg/l; all 11 patients with PGRN-Abs included). IL-1-Ra-Abs were not detected in 28 ICU patients without COVID-19 (Fig. 1 G). Within the discovery cohort titers of IL-1-Ra-Abs ranged between 1:800 to 1:1600 (Fig. 1 H). IL-1-Ra-Abs belonged to IgM and several IgG subclasses, except IgG4 (Fig. 1 I). The epitope region with the highest affinity was located C-terminal from amino acid 63 (Supplementary Fig 2 A).

In a validation cohort of 64 patients (median age: 61 years, 53 male, 11 female) with critical COVID-19 requiring treatment in the ICU, 32 of 64 (50%, median age 60.5 years) had IL-1-Ra-Abs. Similarly to the discovery cohort, IL-1-Ra-Abs in the validation cohort often belonged to both the IgM class and different IgG subclasses and had comparable titers (Supplementary Fig 2 C and D). In contrast, plasma of 5 of 126 (4.0%) hospitalized patients with predominantly moderate COVID-19 contained IL-1-Ra-Abs (supplementary Fig 2 F). Plasma from 2 of 105 (1.9%) patients with mild and asymptomatic COVID-19 contained IL-1-Ra-Abs at low titers, no IL-1-Ra-Abs were detected in 56 convalescent plasma donors of the CAPSID-trial (supplementary Fig. 2 H), and only 2 of 49 (4.1%) patients with mild courses of COVID-19 from the Institute of Virology, Saarland Medical Center (Fig. 1 J and supplementary Fig 2 I) had low titers of Abs. IL-1-Ra-Abs were not detected in serum or plasma of 40 healthcare-workers after 2 vaccinations against SARS-CoV2 (Supplementary Fig. 2 J). As spontaneously occurring IL-1-Ra-autoantibodies had not been reported before, 188 healthy controls (samples from before 2019) were screened but none had IL-1-Ra-Abs (supplementary Fig 2 K). In summary, 42 of 85 critically ill patients (49.4%) had IL-1-Ra-Abs mostly with high titers, compared to 9 of 134 hospitalized patients with moderate or severe COVID-19 (6.7%) (Fisher exact test: *p* < 0.0001), and 2 of 105 patients with mild or asymptomatic COVID-19 (1.9%) with low titers (Fisher exact test: *p* < 0.0001). None of the 75 patients with long-COVID and of the 188 healthy controls (sampled before 2019), and none of the 40 healthy controls vaccinated against SARS-CoV2 had IL-1-Ra-Abs (Figure 1 K) (Fisher exact test: *p* < 0.0001).

Finally, no autoantibodies against IL-10, IL-18BP, IL-22BP, IL-36-Ra, CD40, serpin-B1, IFN-γ or IFN-ω were detected in hospitalized patients with COVID-19 (supplementary Fig 3). Autoantibodies against IFN-α2 were found in 2 of 46 (4.3%) patients with critical COVID-19. Among the hospitalized patients with moderate, severe or critical COVID-19 (n=237) patients of whom survival had been followed-up (n=200), 37 died (18.5%). Of these, 13 were positive for IL-1-Ra-Abs (35%), 11 were positive for PGRN-Abs (29.7%); all the latter were doublepositive for both IL-1-Ra- and PGRN-Abs. Of the 163 survivors, 28 patients were positive for IL-1-Ra-Abs (17.2%), 29 for PGRN-Abs (17.8%) and 18 patients double-positive for both IL-1-Ra-Abs and PGRN-Abs (11%). Thus, without defining a threshold antibody titer, the RR to die for hospitalized patients seropositive for PGRN-Abs was 1.69 (95% CI: 0.9159 to 3.1269, *p*= 0. 0931). For patients seropositive for IL-1-Ra-Abs the RR was 2.10 (95% CI: 1.1748 to 3.756, *p*= 0.0123), and for patients double-positive for IL-1-Ra-Abs and PGRN-Abs the RR was 2.49 (95% CI: 1.3900 to 4.4773, *p*= 0.0022).

### Atypical antigen isoforms in PGRN- and IL-1-Ra-antibody-positive patients

All 11 PGRN-Ab-positive patients of the discovery cohort with severe to critical COVID-19 (36.7%) had a double band of PGRN in isoelectric focusing (IEF), indicating the presence of an additional more negatively charged isoform of PGRN. In contrast, none of the 19 PGRN-Ab-negative patients of this cohort and none of the 28 controls from ICU without COVID-19 produced this signal (Fig. 2 A). ELISA for the previously described pSer81 PGRN isoform(*37*) demonstrated that this more negatively charged isoform of PGRN was indeed pSer81 PGRN (Fig. 2 B). This pSer81 PGRN isoform detected in all seropositive patients with titers ≥ 1:800, but not in patients with low titers (Supplementary Fig. 1 G-H) Antibodies reactive against enriched pSer81 PGRN belonged exclusively to the IgG class, with IgG1 predominating (Supplementary Fig. 4 A). Likewise, in the validation cohort of 64 patients with critical COVID-19, all 25 patients with PGRN-Abs had the hyperphosphorylated PGRN isoform. In non-critical hospitalized patients with antibody-titers below 1:400, none had pSer81 PGRN (Supplementary Fig. 1 E). Some patients developed PGRN-Abs in the further course of disease (Supplementary Fig. 1 F-H).

**Figure 2:**
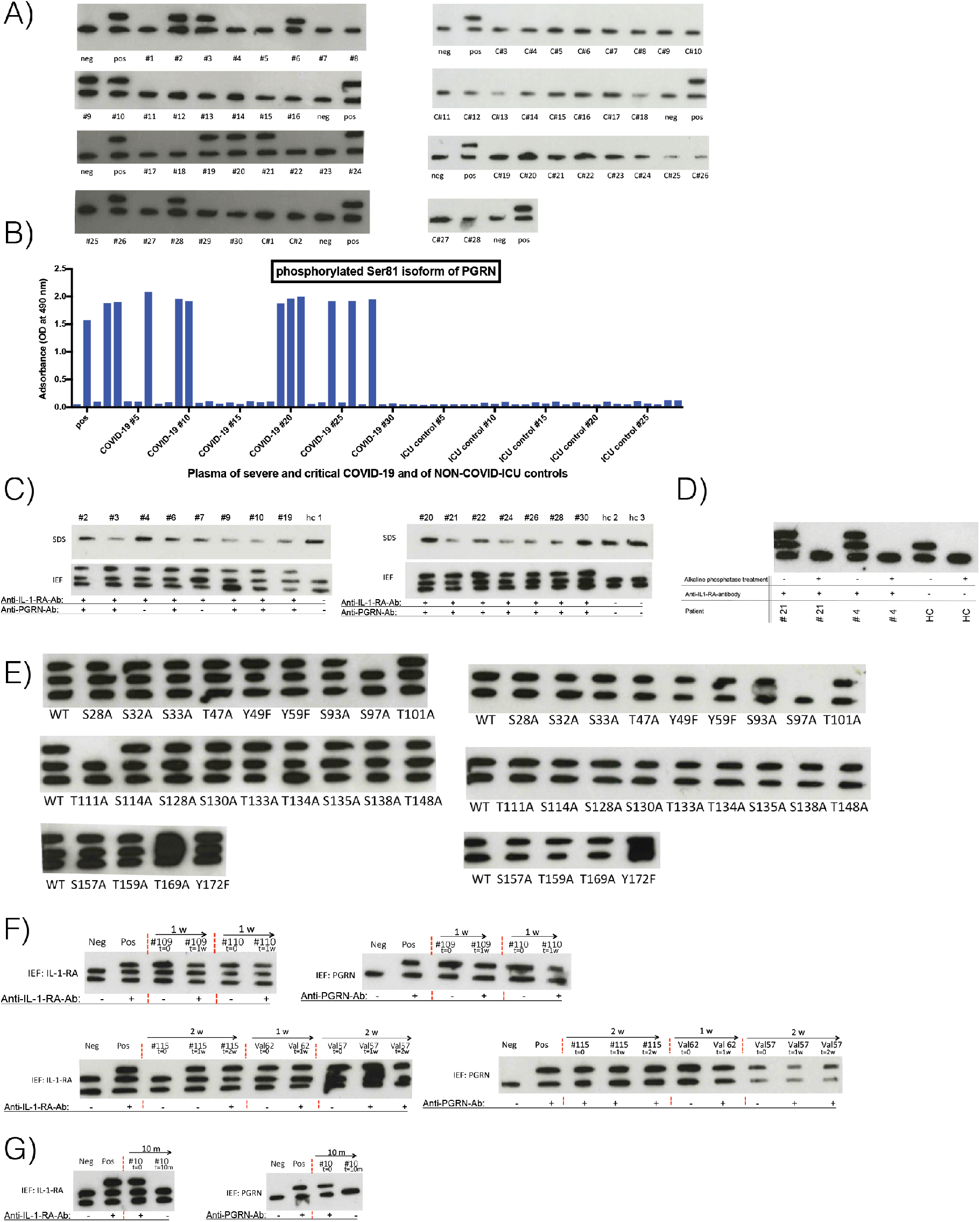
A) IEF of PGRN in plasma from 30 (#1-#30) patients of the discovery cohort with moderate to severe COVID-19 and in 28 controls from patients from ICU without COVID-19. In the samples of patients with PRGN-Abs, an additional and more negatively charged PGRN isoform appeared in the gel, which was not detected in healthy controls and PGRN-Ab-negative patients. B) ELISA for pSer81 PGRN isoform and for non-phosphorylated Ser81 PGRN isoform. C) WB and IEF of IL-1-Ra in IL-1-Ra-Ab-positive patients with moderate to severe COVID-19. An additional and differentially charged IL-1-Ra isoform appeared in patients of the discovery cohort, which was not detected in a healthy controls and IL-1-Ra-Ab-negative patients. D) IEF of IL-1-Ra isoforms from two IL-1-Ra-Ab-positive patients (#21 and #4) and a healthy control without IL-1-Ra-Abs before and after alkaline phosphatase treatment. Alkaline phosphatase treatment led to disappearance of the second phosphorylated isoform, which occurs also in the healthy control, and of the third, atypical and hyperphosphorylated isoform. E) IEF of point-mutated, FLAG-tagged constructs of IL-1-Ra expressed in patient-derived monocytes (left). Only Ser97Ala and Thr111Ala led to the disappearance of the third band of IL-1-Ra. Regarding point-mutated, FLAG-tagged constructs of IL-1-Ra expressed in healthy control-derived monocytes (right) for every construct only two bands were seen, with the exception of Ser97Ala, which led to disappearance of the second band of IL-1-Ra. This indicated Ser97 as physiologic phosphorylation site and Thr111 as pathologic hyperphosphorylation site of IL-1-Ra. F) IEF of IL-1-Ra and PGRN of longitudinally collected samples showed the appearance of the hyperphosphorylated isoforms before antibody seroconversion. This could be detected for IL-1-Ra-Abs and PGRN-Abs in patient #109, #110, Val62 and Val57. In patient #115 initially hyperphosphorylation of PRGN was observed while no hyperphosphorylation of IL-1-Ra was observed. One week later hyperphosphorylated IL-1-Ra appeared, followed by IL-1-Ra-Abs of IgM class one week thereafter. G) In patient #10, who presented initially with critical COVID-19 and was seropositive for PGRN-Abs, IL-1-Ra-Abs and both hyperphosphorylated isoforms, in a follow-up visit 10 month later neither these autoantibodies nor the hyperphosphorylated isoforms were detected anymore. In a further follow-up 14 months neither these autoantibodies nor the hyperphosphorylated isoforms were detected (not shown).

No visible difference in molecular weight of IL-1-Ra was observed in any of the IL-1-Ra-Ab-positive patients with COVID-19 in conventional Western-blot. However, in the IEF, an additional, more negatively charged third band of IL-1-Ra appeared in samples from IL-1-Ra-Ab-positive patients. This isoform of IL-1-Ra was absent in controls without IL-1-Ra-Abs (Fig. 2 C). We then examined whether this additional isoform was related to a different phosphorylation state. Pretreatment with alkaline phosphatase before IEF led to the disappearance of both the normal second and the atypical additional third IL-1-Ra isoform (Fig. 2 D). This indicated that this additional band indeed represented a hyperphosphorylated IL-1-Ra isoform exclusively present in IL1-Ra-Ab-positive patients. Patients with mostly moderate courses of COVID-19 from level 1 and 2 centers of the state of Saarland and seronegative for IL-1-Ra-Abs did not show that third, more negatively charged, hyperphosphorylated isoform of IL-1-Ra (Supplementary Fig. 2 G). After employing patient-derived monocytes and point-mutated FLAG-tagged constructs of IL-1-Ra in Ser97Ala and Thr111Ala the hyperphosphorylated isoform disappeared (Figure 2 E, left). By comparison with transfected monocytes derived from healthy control Ser97 was eventually identified as the physiologically phosphorylated amino acid (second band) and Thr111 as the hyperphosphorylated (atypical third band) amino acid of IL-1-Ra (Figure 2 E, right). Randomly selected patients with either hyperphosphorylated PGRN or hyperphosphorylated IL-1-Ra, dot not have hyperphosphorylated SLP2, which is a previously described hyperphosphorylated antigen of paraproteins in plasma cell malignancies (*39*),(*40*) and served as control (Supplementary Fig. 4 B) (*39*),(*40*).

### Longitudinal analysis of hyperphosphorylated isoforms of IL-1-Ra and PGRN and seroconversion of IL-1-Ra- and PGRN-antibody status

Of 24 patients hospitalized mostly during the second and third SARS-CoV-2-wave in Germany, two to three longitudinal samples obtained at weekly intervals were available. In seven (#109, #110, #104, #115, Val 55, Val57, Val62) patients who were PGRN- and/or IL-1-Ra-Ab-negative at admission, seroconversion to PGRN- and/or IL-1-Ra-Ab-positivity could be observed. In these patients, the hyperphosphorylated isoforms were detected one to two weeks before the appearance of PGRN- and or IL-1-Ra-autoantibodies (Figure 2 F and supplementary Figure 8). In none of the patients the autoantibody preceded the hyperphosphorylated isoform.

Longterm follow-up plasma samples of only one survivor of critical COVID-19 infection of the first wave with initially high-titered IL-1-Ra- and PGRN-Abs and hyperphosphorylated isoforms of both antigens were available. The samples from 10 and 14 months post COVID no longer contained PGRN- and IL-1-Ra-Abs nor hyperphosphorylated PGRN and IL-1-Ra isoforms (Figure 2 G and supplementary Figure 8). Moreover, in 75 patients visiting the long-COVID-19 outpatient ward, no antibodies against PGRN- and IL-1-Ra were detected (Supplementary Figures 1 and 2).

### No cross-reactivity of absorbed PGRN-antibodies against structural proteins of SARS-CoV-2 or Interleukin-1-Receptor Antagonist

Enriched PGRN-Abs failed to show cross-reactivity in ELISA against recombinant HIS-tagged SARS-CoV-2 S1-, S2-, E- or M-proteins, nor against recombinant human FLAG-tagged IL-1-Ra. In addition, antibodies against SARS-CoV-2 S1-, S2-, or M-proteins as well as against human IL-1-Ra could not be adsorbed by immobilized PGRN, but were detectable in the eluate of samples from patients with severe or critical COVID-19, excluding crossreactivity of PGRN-Abs (Supplementary Fig 5).

### Plasma levels of PGRN and IL-1-Ra and functional influence of PGRN-antibody-serostatus on TNF-α-induced cytotoxicity (MTT assay) and of IL-1-Ra-antibody-serostatus on functional IL-1ß-assay

Using standard ELISA, we observed that PGRN levels were significantly decreased by more than 90% in the plasma of PGRN-Ab-positive patients with COVID-19 (median: 15.12 ng/mL; SEM 3.6 ng/mL), compared to (i) plasma of PGRN-Ab-negative patients with COVID-19 (median 161.23 ng/mL, SEM 48.14 ng/mL) (Mann-Whitney test: p < 0.0001) and (ii) plasma of PGRN-Ab-negative patients from the ICU without COVID-19 (median 206.05 ng/mL; SEM 10.59 ng/mL) (Mann-Whitney test: p < 0.0001) (Fig. 3 A). In plasma from a patient with IL-1-Ra-Abs and PGRN-Abs, but not in plasma from a patient seronegative for IL-1-Ra- and PGRN-Abs, immune complexes of both antigens were detectable (Supplementary Fig. 7). Native Western blots under non-reducing conditions and without SDS showed beside converted, mature granulins at approximately 10kDa, reduced levels of free PGRN at approximately 80kDa in PGRN-Ab-positive patients and in addition immunoglobulin-bound PGRN (Fig. 3B). Next, the TNF-a-antagonizing effects of plasma from patients with critical and severe COVID-19 with or without PGRN-Abs were examined with a functional in vitro assay. For this purpose, WEHI-S cells were incubated with TNF-α and plasma at different dilutions from a selected subset of representative patients. As shown in Fig. 3B, addition of plasma from patients without PGRN-Abs reduced the TNF-α-induced cytotoxic effect on WEHI-S cells at higher dilutions when compared to PGRN-Ab containing plasma of patients with COVID-19. This anti-TNF-α effect was detectable up to a plasma dilution of 1:256 or 1:128 with PGRN-Ab-negative, and –positive samples, respectively (Fig. 3 C). The same effect was observed with PGRN-Abs purified from patient plasma or with rec. PGRN-Abs added at different concentrations (Fig. 3 D)

**Figure 3:**
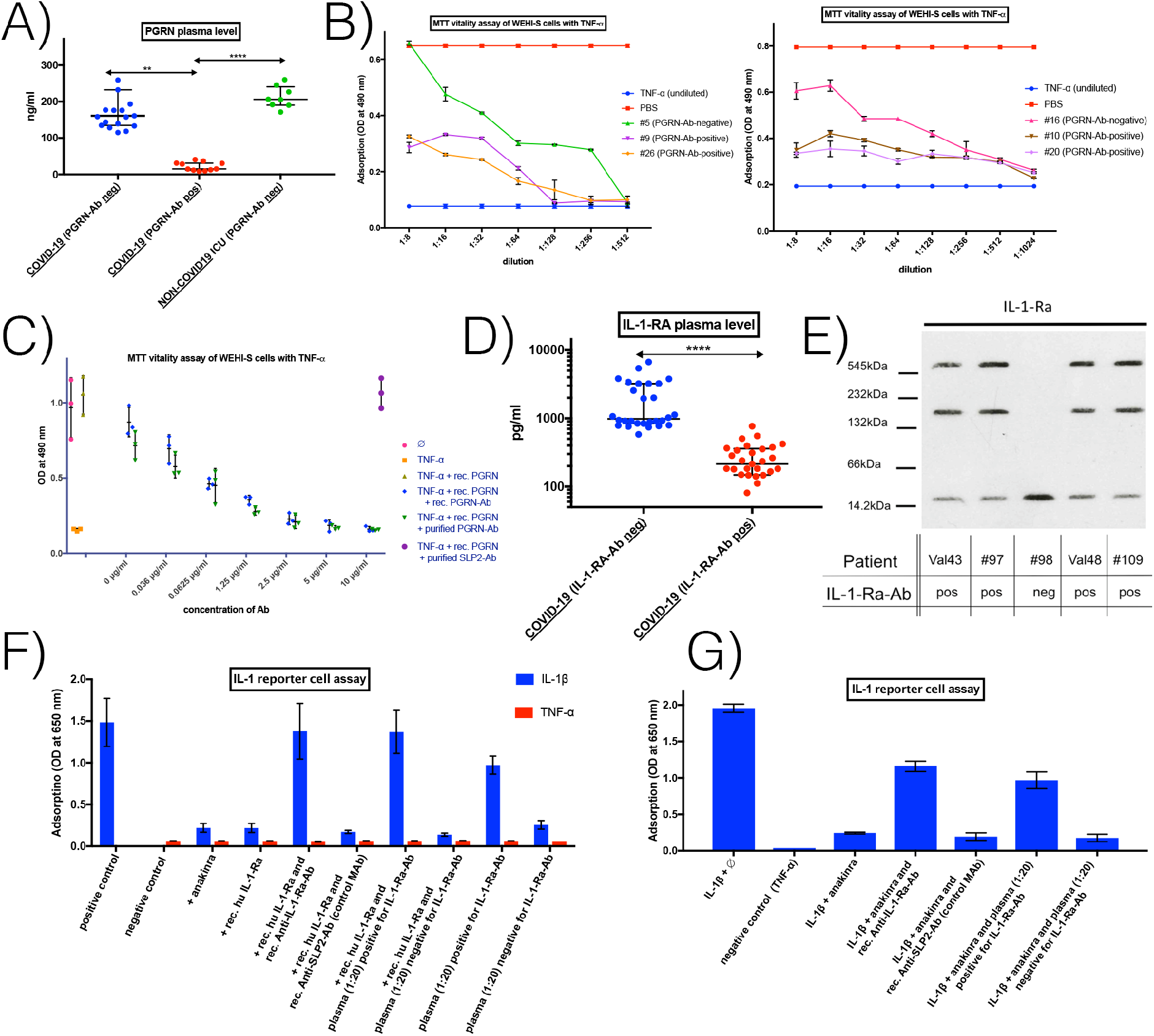
A) PGRN plasma levels in patients with COVID-19 and ICU controls. PGRN plasma levels in controls from ICU without COVID-19 and without PGRN-Abs (median 206.05 ng/ml), and in patients with COVID-19 without PGRN-Abs (median 161.23 ng/ml) were higher compared to high-titer PGRN-Ab-positive patients with moderate to severe COVID-19 (median 15.21 ng/ml). Data are represented with median and interquartile range. B) Native Western blots under non-reducing conditions and without SDS, showed beside converted, mature granulins at approximately 10kDa, reduced levels of free PGRN at approximately 80kDa in PGRN-Ab-positive (Val43, Val48, #109) compared to PGRN-Ab-negative patients with COVID-19 (#97, #98) or with rheumatic disease (rheu1 and rheu2). PGRN-Ab-positive patients (Val43, Val48, #109) had in addition immunoglobulin-bound PGRN, which was not seen for PGRN-Ab-negative patients. C) Effect of PGRN-Ab status on inhibition of TNF-α-induced cytotoxicity. WEHI-S cells were incubated with TNF-α (or PBS) and plasma of PGRN-Ab-positive patients (COVID-19 #9 and #26) or matched PGRN-Ab-negative patient (COVID #5), as indicated. The plasma of the patients #9 and #26 with PGRN-Abs resulted in a weaker inhibition of TNF-α-mediated cytotoxicity. The adsorbance of colored Formazan, as a marker for cell viability was detected at 450 nm. D) Effect of recombinant PGRN-Ab on inhibition of TNF-α-induced cytotoxicity. Purified PGRN-Ab from plasma of patient Val16 both at concentrations from 0μg/ml to 10μg/ml or purified SLP2-Ab at 10μg/ml as a control purified from a patient with multiple myeloma were added to WEHI-S cells incubated with TNF-α and recombinant PGRN. E) IL-1-Ra plasma levels in patients with COVID-19 and ICU controls. IL-1-Ra plasma levels were determined in patients with COVID-19 without IL-Ra-Abs (median 981.8 pg/mL) and in IL-1-Ra-Ab-positive patients with COVID-19 (median 214.4 pg/mL). Data are represented with median and interquartile range. F) Native Western-blot on band strength of IL-1-Ra in IL-1-Ra-Ab-positive or - negative patients. IL-1-Ra-Ab-positive patients had reduced levels of free IL-1-Ra, but additional bands of antibody-bound IL-1-Ra. G) Effect of IL-1-Ra-Ab status on inhibition of IL-1ß. HEK IL-1 reporter cells were incubated with IL-1ß or TNF-α, rec. human IL-1-Ra, anti-IL-1-Ra-Ab, anti-SLP2-Ab as control, and 1:20 diluted plasma of IL-1-Ra-Ab-positive patient (Val 16) or of a matched IL-1-Ra-Ab-negative patient (Val 44), as indicated. The diluted plasma of the patient Val 16 with IL-1-Ra-Abs resulted in a weaker inhibition of IL-1ß. The adsorbance of SEAP, as a marker for IL-1 pathway activation in HEK IL-1 reporter cell was detected at 650 nm. H) Effect of IL-1-Ra-Ab status on inhibition of IL-1ß. HEK IL-1 reporter cells were incubated with IL-1ß or TNF-α, anakinra, anti-IL-1-Ra-Ab, anti-SLP2-Ab as control, and plasma diluted 1:20 of IL-1-Ra-Ab-positive patient (Val 16) or a matched IL-1-Ra-Ab-negative patient (Val 44), as indicated. \*\**P*≤0.01; \*\*\**P* ≤ 0.001; \*\*\*\**P*≤0.0001

Similarly, IL-1-Ra plasma levels were significantly decreased by 78% in IL-1-Ra-Ab-positive patients with COVID-19 (median 214.4 pg/ml; SEM 30.8 pg/ml), compared to plasma of IL-1-Ra-Ab-negative patients with COVID-19 (median 981.8 pg/ml; SEM 301.8 pg/ml) (Mann-Whitney test: p < 0.0001) (Fig. 3E). Native Western-blots showed reduced bands of free IL-1-Ra (18 kDa) in IL-1-Ra-Ab-positive patients compared to IL-1-Ra-negative patients, but IL-1-Ra-Ab-positive patients had in addition immunoglobulin-bound IL-1-Ra (Fig. 3F). An IL-1ß assay was applied to study functional effects of IL-1-Ra-Abs. In HEK IL-1 reporter cells both recombinant human IL-1-Ra and a modified therapeutic version of IL-1-Ra, anakinra antagonized the effect of IL-1ß. Addition of plasma from patients with COVID-19 and seropositive for IL-1-Ra-Abs attenuated the antagonism by recombinant IL-1-Ra or by anakinra, resulting in stronger effects of IL-1ß. This attenuation was comparable to the addition of a recombinant Anti-human IL-1-Ra-Ab (Fig. 3 G and H).

Remarkably, one patient with critical COVID-19 and high titers of PGRN-Abs and IL-1-Ra-Abs required plasmapheresis due to exacerbating chronic inflammatory demyelinating polyneuropathy (CIDP). After 6 days of plasmapheresis, titers of IL-1-Ra and PGRN-Abs strongly decreased from 1:1600 to 1:100, the additional atypic isoform of IL-1-Ra was no longer detectable by IEF in plasma, and the IL-1-Ra plasma levels rose from 81.1 pg/ml to 2340.4 pg/ml (Supplementary Fig. 6).

## DISCUSSION

Here we report the occurrence of autoantibodies to PGRN and IL-1-Ra in a considerable proportion (40% and 49.4%, resp.) of adult patients with critical COVID-19. Importantly, such autoantibodies were not, or barely detectable in COVID-19 patients with moderate, mild or asymptomatic disease, nor in non-COVID-19 ICU patients, SARS-CoV2-vaccinated or in healthy controls. Both IL-1-Ra and PGRN are known to exert anti-inflammatory activity, suggesting a causal link to the hyperinflammatory phenotype often observed in critically ill COVID-19 patients. It is noteworthy that PGRN-autoantibodies were first identified in plasma of 20-40% of patients suffering from primary small, middle and large vessel vasculitis (*23*). Critical forms of COVID-19 have been shown to be frequently associated with vasculitis-like characteristics (*4*), which may contribute to some of the complications of this disease.

IL-1-Ra is the physiologic inhibitor of IL-1α and IL-1β due to competition for IL-1R1. A proinflammatory imbalance between these proteins is found in gout crystallopathies, autoinflammatory or cryopyrin-associated periodic fever syndromes. Macrophage activation syndrome, which has many similarities to HLH, is a common complication in systemic juvenile idiopathic arthritis and adult-onset Still’s disease. These conditions have in common the activation of the NLRP3 inflammasome (*41*). Therapeutically, colchicine, recombinant human IL-1-Ra or monoclonal antibodies against IL-1β are used (*42*),(*10*). To our knowledge, except for IL-1-Ra anti-drug-antibodies (*43*), spontaneously occurring IL-1-Ra-Abs had not been described before in the literature. Shortly after the publication of the preprint of the present manuscript (*44*), neutralizing IL-1-Ra-Abs were reported in IgG4-associated diseases (*45*). Appearance of IL-1-Ra-Abs was predominantly observed in patients with critical COVID-19, barely in patients with mild or asymptomatic COVID-19, not in non-COVID-19 patients at internal medicine ICU, not in healthy controls and not in SARS-CoV-2-vaccinated healthy controls. These antibodies will have to be characterized in more detail and investigation of their potential occurrence in rheumatologic or autoinflammatory disorders beyond IgG4-associated diseases (*45*) may also be worthwhile in future studies.

What might have triggered the production of PGRN- and IL-1-Ra-Abs? A noteworthy observation in this context is that in contrast to rheumatologic diseases, where PGRN-Abs largely belong to the IgG1 class(*23*), in COVID-19 PGRN-Abs were distributed across the IgM and multiple IgG subclasses. Moreover, the PGRN-Ab titers were high, ranging from 1:1600 up to 1:3200 (Fig. 1 C-D). Similarly, IL-1-Ra-Abs were found at high titers and were also distributed across IgM and different IgG subclasses (Fig. 1 H-I). These observations suggest that a recent event might have triggered the formation of these antibodies – possibly the COVID-19 infection itself. No cross-reactivity of PGRN-Abs with SARS-CoV-2 structural proteins was observed. Atypical posttranslational modifications may represent a common mechanism provoking autoimmunity (*46*),(*47*),(*40*),(*48*). In patients with severe to critical COVID-19 with relevant titers of PGRN-Abs (≥1:800), the pSer81 PGRN isoform was consistently found (in addition to the non-phosphorylated isoform), whereas pSer81 PGRN was not detected in PGRN-Ab-negative subjects. In previous studies on autoimmune diseases, we had identified a posttranslationally modified pSer81 PGRN isoform exclusively in PGRN-Ab-positive patients, preceding antibody formation, as the reason for the immunogenicity of PGRN (*37*). Moreover, we found that this second pSer81 isoform led to altered functions of PGRN, with a dramatic reduction in its affinity to TNFR1/2 and DR3 and consequently, a loss of PGRN’s ability to antagonize the TNF-α and TL1A effects (*37*). In autoimmune diseases PKCß1 was identified as the kinase and PP1 as the phosphatase relevant for phosphorylation and dephosphorylation of Ser81 PGRN, and inactivation of PP1 seemed to be responsible for the observed phosphorylation of PGRN at the Ser81 residue (*37*).

Similarly, in our present work, we found that IL-1-Ra presented as an additional, hyperphosphorylated isoform. It was present exclusively in all patients with high titers of antibodies against IL-1-Ra. Ser97 was identified as a physiologically phosphorylated site and Thr111 as the atypically hyperphosphorylated amino acid of IL-1-Ra.

Interestingly, similar to the reported preceding appearance of pSer81 PGRN prior to PGRN-Abs in rheumatic diseases (*37*), in COVID-19 we observed this phenomenon of seroconversions for IL-1-Ra- and PGRN-Abs following the initial appearance of hyperphosphorylated isoforms of both IL-1-Ra and PGRN in 7 of 24 hospitalized patients with available longitudinal samples. These observations suggest that the posttranslational immunogenic modifications of PRGN and IL-1-Ra are induced by the SARS-CoV-2-infection itself or the inflammatory environment evoked by the infection.

Further studies will be needed to clarify the processes underlying this autoimmune response to IL-1-Ra and PGRN in COVID-19, for instance a screening of inflammatory mediators as potential triggers for the hyperphosphorylation of PGRN and IL-1-Ra might help to identify susceptible individuals, and the identification of the responsible kinase(s) and phosphatase(s), as well as characterization of T-helper cell responses against hyperphosphorylated epitopes of IL-1-Ra and PGRN might provide a deeper understanding.

Whatever the primary cause of the loss of self-tolerance to PGRN and IL-1-Ra may be, the present observations raise the question what potential clinical implications the autoantibodies might have. Notably, PGRN and IL-1-Ra plasma levels were substantially reduced in patients with PGRN-Abs and IL-1-Ra-Abs, respectively, as compared to seronegative patients (Fig 3 A). This contrasts with the finding that patients with COVID-19 had elevated PGRN and IL-1-Ra plasma levels, as detected by multiplex assays (*49*)(*50*)(*51*)(*52*). Native Western blots could possibly provide an explanation for this discrepancy. Levels of free IL-1-Ra and PGRN were strongly decreased, and Ig-bound IL-1-Ra and PGRN represented the majority in antibody-positive patients (Fig. 3 B and 3 F). Dilution series with rec. IL-1-Ra with or without addition of rec. or purified IL-1-Ra-Abs validated the commercial ELISA and showed, that it measures only free IL-1-Ra (supplementary Fig. 9).

Functional confirmation of the neutralizing effect of PGRN-Abs was obtained in a TNF-α-induced cytotoxicity assay, which showed that plasma from PGRN-Ab-positive patients was less effective than plasma from PGRN-Ab-negative patients in inhibiting the effect of TNFa in vitro (Fig. 3 C and 3 D). The proinflammatory capacity of IL-1-Ra-Abs was confirmed with HEK IL-1 reporter cells, in which IL-1-Ra-Abs functionally neutralized both recombinant human IL-1-Ra and anakinra (Fig. 3 G and H). It thus appears plausible that the autoantibodies detected in the present study have caused the observed massive reduction in the circulating levels of PGRN and IL-1-Ra. This notion is further supported by the observation of immunocomplexes of PGRN and IL-1-Ra. It is thus tempting to speculate that this reduction of two anti-inflammatory regulators might contribute to a proinflammatory milieu in a relevant subgroup of critically affected patients with COVID-19 and is associated with significantly higher morbidity and mortality. The resulting proinflammatory shift in the inflammatory balance due to PGRN-Abs and IL-1-Ra-Abs would represent a different mechanism compared to autoantibodies neutralizing type I IFN, which primarily weaken the antiviral immune response(*16*). In addition to the in vitro data, PGRN- and IL-1-Ra-Abs were significantly more frequently observed in patients who required intensive care treatment. Furthermore, the presence of IL-1-Ra-Abs or of IL-1-Ra-Abs in combination with PGRN-Abs was associated with an increased mortality from COVID-19 in hospitalized patients.

The current results raise several intriguing questions for future investigations. In particular, it will be interesting to clarify i) who is susceptible to develop hyperphosphorylated immunogenic isoforms of PGRN and IL-1-Ra and ii) what are the molecular underlying mechanisms for this formation of hyperphosphorylated isoforms in the context of COVID-19. Another question is whether the presence of PRGN-Abs and IL-1-Ra-Abs (and the ensuing neutralization of PGRN and IL-1-Ra) in patients with critical COVID-19 represents a causal factor in the development of an autoimmune-like vasculitis (*4*).

A further potential implication of autoantibody-induced downregulation of two key inflammatory pathways is that targeted therapeutic approaches for this subgroup of patients might consist in targeted reinforcement of these impaired anti-inflammatory pathways.

A possible next step towards evaluating the potential of a more-targeted therapy might be to perform post-hoc analyses of plasma samples from prospective trials investigating agents modulating these pathways and related pathways like IL-6 and to look for correlations with baseline characteristics, response and outcome (*6*),(*9*),(*8*),(*7*),(*53*),(*11*),(*54*). This would be of particular interest for the modulation of the IL-1 pathway, as anakinra, which is a recombinant fragment spanning from amino acids 26 to 177 of human IL-1-Ra, was in-vitro functionally neutralized by plasma of IL-1-Ra-Ab-positive patients. However, it is not possible to draw a simple conclusion about the clinical effect of anakinra in IL-1-Ra-Ab-positive patients. Furthermore, the presence of IL-1-Ra-Abs has to be investigated in other autoimmune and autoinflammatory diseases.

Our study has several limitations. Unfortunately, no samples from prospective clinical trials or from different geographic regions and ethnic groups could be examined, and. Another weakness is that only selected known proinflammatory autoantibodies as PGRN- or IL-10-Abs or hypothesis-based possible candidate autoantibodies were investigated. This limited search, on the other hand, has led to a focused and detailed characterization.

In addition, the previously unknown hyperphosphorylated isoform of IL-1-Ra should be further characterized in future studies, similar to the reported distinct function of pSer81 PGRN in autoimmune diseases (*37*). Future studies need to verify the frequency of these autoantibodies in critical COVID-19, to analyze the time course of their occurrence and their clinical relevance including the prognostic and predictive value and therapeutic options. Finally, it should be addressed, whether these or other pathogenic autoantibodies play a role in multisystem inflammatory syndrome in children (MIS-C).

## MATERIAL AND METHODS

This study was approved by the local Ethical Review Board (Bu 62/20) and conducted according to the Declaration of Helsinki. Plasma samples and PBMCs from a discovery cohort of 30 adult patients with critical (n=21) or severe (n=9) COVID-19 of the first wave of the pandemic in the spring of 2020 and from 28 patients without COVID-19 requiring intensive care treatment were obtained from the COVID-19 wards of the Department of Internal Medicine II and V and the ICU of the Department of Internal Medicine III of the Saarland University Hospital (Homburg/Saar, Germany) and were analyzed in an extended in house diagnostics for proinflammatory autoimmunity.

For validation, blood samples from 64 adult patients with critical, life-threatening COVID-19 requiring mechanical ventilation and/or ECMO treatment of the Department of Internal Medicine V of the Saarland University Hospital (Homburg/Saar, Germany) and of patients treated within the CAPSID trial (EUDRA-CT 2020-001310-38) were analyzed for proinflammatory autoimmunity. Plasma samples and whole blood lysates from a control group of adult hospitalized patients (n=125) with predominantly moderate COVID-19 at the time of admission and blood sample collection were obtained from level 1 and 2 hospitals in Saarland federal German state.

Regarding asymptomatic to mild courses of COVID-19, plasma and serum samples of 105 adult patients were obtained, 49 were provided by the Institute of Virology of the Saarland Medical Center Homburg/Germany and 56 originated from convalescent plasma donors of the CAPSID trial (EUDRA-CT 2020-001310-38) from the Institute of Clinical Transfusion Medicine and Immunogenetics at the University Hospital Ulm (Germany). Blood samples of 40 vaccinated healthcare workers were obtained from MVZ Mindelheim/Germany and from the Institute of virology of Saarland Medical Center Homburg/Germany. Moreover, plasma samples from 75 patients were obtained from the long-COVID outpatient ward of the Department of Internal Medicine V of the Saarland University Hospital (Homburg/Saar, Germany). Finally, plasma and whole blood samples from 188 adult healthy controls were obtained before 2019. All samples were from adult patients. All plasma and PBMC samples were stored at - 20°C or - 80°C, respectively until use.

### ELISA for autoantibodies against PGRN, IL-1-RA and IL-10, IL-18BP, IL-22BP, IL-36-Ra, CD40, serpinB1, IFN-α2, IFN-γ and IFN-ω

The ELISA for autoantibodies was performed as previously described (*55*). In short, the antigens were obtained using the coding sequences of the *GRN* gene encoding PGRN, IL-10 and isoform 1 precursor and isoform 2 of *IL1RN*, and the fragments aa1-63, aa107-177 and aa47-123 of isoform 1 of IL-1-Ra, IL-18BP, IL-22BP, IL-36-Ra, CD40, serpinB1, IFN-α2, IFN-γ and IFN-ω which were recombinantly expressed with a C-terminal FLAG-tag in HEK293 cells under the control of a cytomegalovirus promoter (pSFI). Total cell extracts were prepared and bound to Nunc MaxiSorp plates (eBioscience, Frankfurt, Germany) precoated with murine anti-FLAG mAb at a dilution of 1:2,500 (v/v; Sigma-Aldrich, Munich, Germany) at 4°C overnight. After blocking with 1.5% (w/v) gelatin in Tris-buffered saline (TBS) and washing steps with TBS with Triton X-100, the individual plasma samples were diluted 1:100. ELISA was performed according to standard protocols with the following Abs: biotinylated goat antihuman heavy and light chain immunoglobulin G (IgG) at a dilution of 1:2,500 (Dianova, Hamburg, Germany); subclass-specific sheep antihuman IgG1, IgG2, IgG3 and IgG4 (Binding Site Group, Birmingham, UK) at dilutions of 1:5,000; goat antihuman IgM (Dianova) at a dilution of 1:2,500; or goat antihuman IgA (Dianova) at a dilution of 1:2,500. Following this step, corresponding biotinylated secondary Abs were used for immunoassays carried out to detect IgG subclasses and IgM. Peroxidase-labelled streptavidin (Roche Applied Science, Indianapolis, IN, USA) was used at a dilution of 1:50,000. As a cut-off for positivity, the average of the optical density (OD) of the negative samples plus three standard deviations was applied. For the determination of the binding region of IL-1-Ra, recombinant fragments of different lengths were constructed with C-terminal FLAG tags and expressed as described earlier.

### Analysis of IL-1-Ra- or PGRN-immunocomplexes

To investigate the presence of immunocomplexes of PGRN or IL-1-Ra, rabbit anti human IL-1-Ra-Abs (antibodies-online # ABIN2856394) or mouse anti-human PGRN-Abs (abcam#ab169325) at 5 μg/mL were incubated and coupled with Affi-gel Hz Hydrazide (Biorad #156-0016). Plasma 1:100 diluted from a patient with critical COVID-19 (Val 51) seropositive for PGRN-Abs and IL-1-Ra-Abs and a patient (Val 52) with critical COVID-19 seronegative for PGRN-Abs and IL-1-Ra-Abs were incubated at 4 °C overnight with either Affi-matrix with rabbit anti human IL-1-Ra-Abs or with mouse anti-human PGRN at 5 μg/mL. Elution was done with 0.25M glycine pH 3. Purified products were separated by SDS-PAGE and after blocking for Western Blot either murine PGRN antibody (abcam#ab169325) followed by anti-mouse/POX 1:3000 (Biorad#170-6516), rabbit IL-1-Ra antibody (antibodies-online # ABIN2856394) followed by anti rabbit/POX 1:3000 (Biorad#170-6515), or by biotinylated anti-human IgG antibody (Dianova#109-066-097) followed by Strep/Pox 1:15000 (Roche#11089153001) were used.

### Western blot, isoelectric focusing of PGRN and IL-1-Ra, and ELISA for pSer81 or npSer81 PGRN

Western blotting and isoelectric focusing was performed as described (*37*). Whole blood lysates or lysates of PBMCs from PGRN- and/or IL-1-Ra-Ab-positive patients and controls were analyzed by IEF for PGRN and SLP2 isoforms and plasma samples were analyzed for IL-1-Ra isoforms. Whole blood cell lysates or lysates of PBMCs from IL-1-Ra-Ab positive patients were treated with alkaline phosphatase as previously described using FastAP thermosensitive alkaline phosphatase (Fermentas/VWR, Darmstadt, Germany).(*40*) In addition for detection of free or Ig-bound IL-1-Ra or PGRN Western-blots without reducing sample preparations and without SDS were performed. For ELISA for the pSer81 isoform of PGRN Nunc MaxiSorb plates were coated overnight at 4°C with rabbit antihuman PGRN antibodies directed against the C-terminus at a dilution of 1:2500 (v/v; LsBio, Seattle,WA, USA), followed by blocking with 1.5% (w/v) gelatin in TBS and washing steps with TBS-Tx [TBS, 0.1% (v/v) Tx100]. Individual plasma samples were utilized at a dilution of 1:2, and individual whole blood cell lysates were utilized at a dilution of 1:100. ELISA was performed according to standard protocols. For the detection of the hyperphosphorylated pSer81 or the non-phosphorylated Ser81 PGRN isoform, phospho-Ser81- or non-phospho-Ser81 PGRN-specific Fabs, which had previously been selected by phage display screening (*37*), were used at a concentration of 10 μg/ml. Following this, corresponding biotinylated antihuman Fab secondary antibodies and subsequently peroxidase-labeled streptavidin (Roche) were used.

### Site-directed mutagenesis

Using QuickChange II Site-Directed Mutagenesis Kit (Stratagene) and *IL1RN* full-length FLAG-tagged DNA fragments, all candidate phosphorylation sites were mutagenized. In each case, the amino acids serine (S) and threonine (T) were replaced by alanine (A) and tyrosine (Y) was replaced by phenylalanine (F) disabling phosphorylation. This exchange was performed for *IL1RN* with S28A, S32A, S33A, T47A, Y49F, Y59F, S93A, S97A, T101A, T111A, S114A, S128A, S130A, T133A, T134A, S135A, S138A, T148A, S157A, T159A, T169A and Y172F. These FLAG-tagged mutants were cloned into pRTS and expressed in electroporated primary monocytes isolated by CD14 MACS (Pan Monocyte isolation kit, Miltenyi, Germany) from of a patient with hyperphosphorylated IL-1-Ra and IL-1-Ra-Abs and a control without hyperphosphorylated IL-1-Ra and negative for IL-1-Ra-Abs and subsequent cultivation with RPMI and 20% FCS with M-CSF at 10ng/ml and IL-4 at 1ng/ml.

### ELISA for antibodies against pSer81 PGRN isoform and their Ig class

To detect antibodies against pSer81 PGRN, Nunc MaxiSorp plates (eBioscience, Frankfurt, Germany) were precoated with murine anti-HIS mAb at a dilution of 1:2,500 (v/v; Sigma-Aldrich, Munich, Germany) at 4°C overnight. After blocking with 1.5% (w/v) gelatin in Trisbuffered saline (TBS) and washing steps with TBS with Triton X-100, HIS-tagged pSer81-specific recombinant Fabs (which had previously been selected by phage display screening (*37*)), were added at 10 μg/ml followed by washing steps with TBS with Triton X-100 and addition of lysates of PBMCs of patients with the pSer81 isoform. This was followed again by washing steps with TBS with Triton X-100 and by addition of individual plasma samples were diluted 1:100. Biotinylated goat antihuman heavy and light chain IgG were used at a dilution of 1:2,500 (Dianova, Hamburg, Germany); subclass-specific sheep antihuman IgG1, IgG2, IgG3 and IgG4 (Binding Site Group, Birmingham, UK) at dilutions of 1:5,000; and goat antihuman IgM (Dianova) at a dilution of 1:2,500. Following this step, corresponding biotinylated secondary Abs were used for immunoassays carried out to detect IgG subclasses and IgM. Peroxidase-labelled streptavidin (Roche Applied Science, Indianapolis, IN, USA) was used at a dilution of 1:50,000.

### ELISA for plasma level determination of PGRN and IL-1-Ra

PGRN plasma levels were determined of 19 patients with COVID-19 without PGRN-Abs, 11 patients with COVID-19 with PGRN-Abs and 8 patients without COVID-19 and without PGRN-Abs treated on ICU with a commercially available ELISA kit (AdipoGen, Incheon, South Korea) according to the manufacturer’s instructions. IL-1-Ra plasma levels were determined of 20 patients positive and 12 patients negative for IL-1-Ra-Abs with a commercially available ELISA kit (Invitrogen/ThermoFisher #BMS2080) according to the manufacturer’s instructions. To investigate if the commercial ELISA-kits detect only free IL-1-Ra and PGRN, or also the Ig-bound IL-1-Ra and PGRN, a dilution series with recombinant IL-1-Ra or PGRN were performed from 10,000 pg/ml, 1000 pg/ml, 100 pg/ml, 10 pg/ml and 0 pg/ml with or without the addition of 0.05μg rec. IL-1-Ra-Abs or rec. PGRN-Abs or from patient’s plasma (Val16) purified IL-1-Ra-Abs or PGRN-Abs.

### TNF-α-induced cytotoxic effect (MTT assay)

To assess the functional effects of PGRN-Abs in vitro, a nonradioactive viability assay (EZ4U Cell Proliferation Assay; Biomedica, Vienna, Austria) was performed. For this TNF-α-induced cytotoxicity indicator assay, we used the highly TNF-α-sensitive mouse fibrosarcoma WEHI-S cell line as target cells. In short, 4×10^4^ WEHI-S cells were seeded into 200 μl of cell culture at 37°C and 5% CO2. To detect possible differences of TNF-α inhibiting activity in plasma between patients with or without PGRN-Abs, plasma of patients with COVID-19 with or without PGRN-Abs was added in dilutions from 1:8 to 1:512 to cultured WEHI-S cells, followed by administration of TNF-α (100 pg/ml). WEHI-S cells without addition of TNF-α and plasma, or solely with addition of TNF-α (100 pg/ml), were used as positive and negative controls, respectively. After 48 hours of incubation at 37°C, chromophore substrate was added to each well. This chromophore substrate is converted only by vital cells. The adsorption of the product was measured at an OD of 450 nm. Nonetheless the very short halflive of TNF-α compared to PGRN, to exclude any possible unknown interferences from other plasma components, the MTT assay was repeated with TNF-α (100 pg/ml) and recombinant PGRN (10ng/ml) with either recombinant PGRN-Abs, PGRN-Abs purified from plasma of patient Val16 at concentrations from 0μg/ml to 10 μg/ml or as a control SLP2-paraprotein purified from a patient with multiple myeloma at 10 μg/ml(*56*).

### IL-1ß-assay

For IL-1ß assay HEK-Blue^™^ IL-1β reporter cells (Invivogen, #hkb-il1bv2) were used, which react specifically to IL-1ß and IL-1α by induction of NF-κB/AP-1, leading to expression of a secreted embryonic alkaline phosphatase (SEAP) reporter. Anakinra at 40ng/mL, recombinant IL-1-Ra at 40ng/mL (Biozol, #PPT-AF-2000-01RA), recombinant IL-1-Ra at 40ng/mL (Biozol, #PPT-AF-2000-01RA) and anti-IL-1-Ra antibody at 5 μg/mL (antibodies-online#ABIN2856394), recombinant IL-1-Ra at 40ng/mL (Biozol, #PPT-AF-2000-01RA ml and recombinant SLP-antibody at 5μg/mL (abcam, #ab191883), plasma diluted 1:20 of a patient with COVID-19 and high-titered IL-1-Ra-antibodies (Val 16) with and without recombinant IL-1-Ra at 40ng/mL, and plasma diluted 1:20 of a patient with COVID-19 without high-titered IL-1-Ra-Abs (Val 44) with or without recombinant IL-1-Ra at 40ng/mL were preincubated for 2h at room temperature. Subsequently, these compounds were added with either IL-1ß (Biozol,# PPT-200-01B) or TNF-α (Biozol, #PPT-300-01A) both at 2ng/mL in 100μl DMEM to 2×10E4 HEK-Blue^™^ IL-1β reporter cells per well and incubated overnight at 37°C. Thereafter, 180 μL of each supernatant was transferred, 20μl QUANTI-BlueTM (Invivogen, #rep-qbs) was added and SEAP activity was measured at OD of 655nm. Experiments were performed in triplicate.

### ELISA for cross-reactivity of absorbed PGRN-antibodies against structural proteins of SARS-CoV-2

PGRN-Abs were enriched using plasma from patients listed below. For this purpose, 500 μl of lysate of HEK293 cells transfected with recombinant FLAG-tagged PGRN was incubated with 20 ml anti-FLAG matrix for 15 min at room temperature. PGRN-Ab positive patient’s plasma (500 μl) was diluted 1:10 (v/v) in PBS and was incubated with the anti-FLAG matrix/FLAG-tagged PGRN complex and subsequently desorbed by elution with glycine buffer depleted. Elution was performed with 100 ml of 0.1 M glycine pH 3. The patient plasma, the flow-through, the eluted enriched PGRN-Abs and plasma of controls listed below were screened by ELISA for reactivity against recombinant HIS-tagged SARS-CoV-2 S1- and S2-proteins, N-protein and M-protein expressed in HEK293 cell (ABIN) and against reactivity against FLAG-tagged PGRN, precursor of IL-1-Ra isoform 1 and IL-1-Ra isoform 2. PGRN-Abs were purified from plasma patient #10 and #20 of the cohort with moderate and severe COVID-19 infection, respectively. Plasma from two healthy control, from a patient with rheumatologic disease with PGRN-Abs and without COVID-19, from a patient with rheumatologic disease without PGRN-Abs and without COVID-19 served as controls.

### Data analysis

Comparrison of proportions was done with two-tailed Fisher exact test. IL-1 Ra and PGRN plasma levels were analyzed with D’Agostino and Pearson normality test and Shapiro-Wilk normality test. Plasma levels were then compared by nonparametric, two-tailed Mann-Whitney test.

**Table 1:**
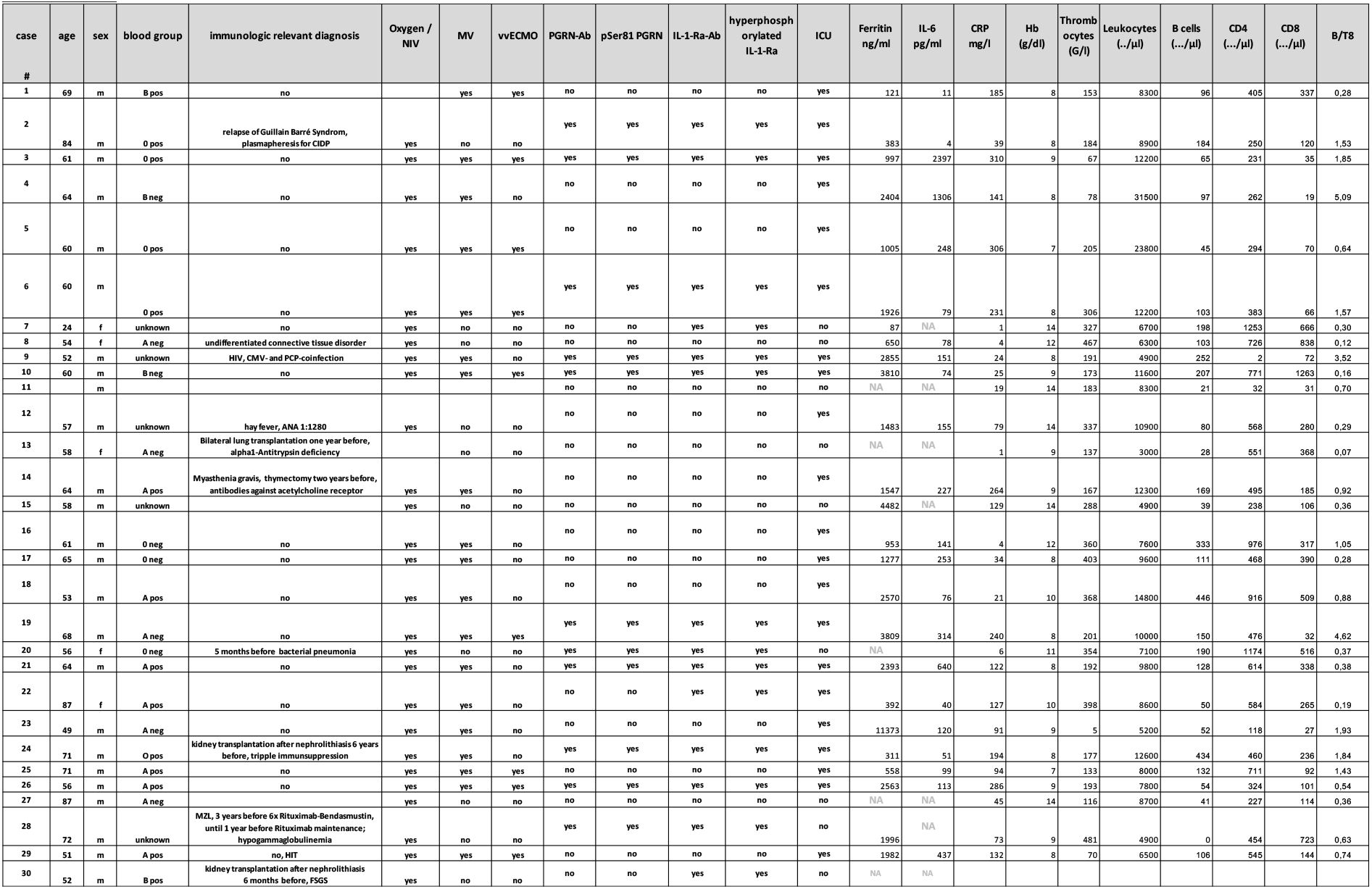
Patients’ characteristics and antibody-status of discovery cohort of 30 patients with severe or moderate COVID-19

## Author contribution

LTh, NF, ER and BTh planned the study. NF and ER performed experiments. LTh, BTh and YF wrote the manuscript. LTh, BTh, YF, NF, ER, KDP, SH, DKM, TR, CP, GG, JP, SB, MB, MR, KC, AB, VL, MA, MB, SS, FL, RB, SB, SM, MK and PL revised the data and manuscript. IK, MB, BTh, CS, FN, JP, TR, SM, AB, PA, CM. FS, JL, TA, SE, AL, CW, AT, MR, BF, GD, CP, TP, JR, MB, MH, RB, FL, SL, SM, CH, CL, KR, SH, SS, SB, MK, PL, SK and HS provided samples of patients and controls and clinical data. FL, GG and BF contributed excellent medical assistance without which, the conduct of the study would not have been possible.

## Acknowledgement

This work was supported by a young investigator NanoBioMed fund of the University of Saarland to LTh. This work was supported by the CorSAAR-registry. The work was partly supported by the Ministry of Science, Research and the Arts of the State of Baden-Württemberg within the special funding initiative COVID-19 research (“CORE-Study – COVID-19 adult reconvalescent plasma”).

